# A Stretchable Wearable Doppler Ultrasound Patch for Continuous Vascular Monitoring

**DOI:** 10.1101/2025.11.23.690069

**Authors:** Yuanlong Li, Ziqi Li, Chang Peng

**Affiliations:** School of Biomedical Engineering & State Key Laboratory of Advanced Medical Materials and Devices, ShanghaiTech University, Shanghai, 201210, China

**Keywords:** Blood flow monitoring, wearable ultrasound, Doppler ultrasound, slow-time sampling

## Abstract

Continuous blood flow monitoring is crucial for the diagnosing and preventing cardiovascular diseases, but existing techniques have several limitations that hinder widespread clinical use, such as invasiveness, operator dependency and lack of continuous measurement. In this paper, we present a wearable ultrasound patch designed for continuous blood flow monitoring. The patch integrates six ultrasonic transducers arranged symmetrically along the central axis. A baseplate with a specific angle (30 degrees) is used to form a 60-degree Doppler incident angle. We have simplified and improved the fabrication process of the serpentine electrodes, avoiding the cumbersome traditional steps such as transfer printing and spin-coating of Cu/PI. The fabricated wearable ultrasound patch exhibits a center frequency of 5.05 MHz, a -6 dB bandwidth of 42.58%, and an elastic stretchability of up to 30%. Based on pulsed-wave (PW) Doppler, it calculates the Doppler frequency shift using a slow-time sampling algorithm. In vitro studies showed an absolute percentage error of 4.1% to 12% for constant flow velocity measurements, with actual flow velocities varying between 20 and 100 cm/s. The mean absolute error (MAE) was 3.56 cm/s, and the mean absolute percentage error (MAPE) was 5.99%. In pulsatile flow experiments, the patch was also able to capture the changing peaks of the velocity waveform, demonstrating its feasibility for continuous blood flow monitoring.

## 1. Introduction

Cardiovascular disease is the leading cause of death worldwide [1]. According to data from the World Health Organization (WHO), approximately 17.9 to 19 million people die from cardiovascular diseases each year, accounting for more than 31% of all global deaths [2, 3]. In China, cardiovascular and cerebrovascular diseases have become the "number one killer" of public health, with two out of every five deaths attributed to these diseases, representing 40% of all disease-related mortality [4]. Hemodynamics, a discipline focused on studying the mechanical principles of blood flow within the cardiovascular system, provides indispensable scientific support for investigating the pathogenesis of cardiovascular diseases, evaluating clinical conditions, formulating treatment strategies, and assessing prognosis [5, 6, 7]. This is achieved through precise measurement and analysis of key parameters such as blood pressure, blood flow, vascular resistance, and blood flow velocity. Among various hemodynamic parameters, blood flow velocity holds a central role due to its ability to directly and sensitively reflect vascular functional status, tissue perfusion efficiency, and disease severity. Changes in blood flow velocity often serve as early indicators of initial vascular pathologies, such as endothelial dysfunction or microcirculation impairment, preceding many structural changes. In the assessment of vascular stenosis, peak systolic velocity and end-diastolic velocity have become the clinical diagnostic "gold standard" [8, 9, 10]. Therefore, as a dynamic and comprehensive parameter rich in pathophysiological information, the accurate assessment of blood flow velocity is essential for the comprehensive management of cardiovascular diseases.

Currently, clinical methods for blood flow monitoring can be clearly categorized as invasive and non-invasive approaches [11]. Among them, invasive methods require the surgical placement of a catheter or sensor directly into human blood vessels. By virtue of “direct contact with blood flow,” these techniques are often regarded as the “gold standard” for diagnosis in specific clinical scenarios, offering not only high accuracy but also real-time monitoring capabilities [12]. However, such methods have significant limitations: the insertion procedure increases the risk of vascular infection and may cause patient discomfort or other complication [13]. More importantly, due to their invasive nature, they are unsuitable for long-term or continuous blood flow monitoring, which greatly restricts their application in the prolonged management of patients with chronic conditions. In contrast, non-invasive monitoring methods, which preserve vascular integrity and are convenient to operate, have become the preferred choice in a growing number of clinical settings [14, 15]. Among non-invasive techniques, ultrasonic Doppler, magnetic resonance angiography (MRA), and laser Doppler flowmetry are three classical types. While MRA can clearly visualize blood flow distribution, it involves high equipment costs, expensive procedures, and lengthy scanning times [16]. Laser Doppler flowmetry, limited by its technical principles, has a shallow penetration depth (only 2–4 cm) and is unable to effectively assess blood flow in deep human vessels or organ [17]. Doppler ultrasound technology, with operational convenience, real-time monitoring, and high efficiency, is widely used in clinical hemodynamic assessments such as peripheral vascular screening and cardiac blood flow evaluation [18, 19]. However, most conventional Doppler ultrasound probes are rigid, relatively bulky, and difficult to conform stably to the skin. Their use requires the application of hydrogel as a coupling agent, which tends to dry out or dislodge, making them unsuitable for wearable continuous monitoring [20]-[22]. Furthermore, the measurement process relies on manual operator adjustment for probe angle and pressure, greatly limiting their applicability in home-based monitoring or emergency bedside scenario [23, 24, 25].

In recent years, the rapid development of flexible electronics has brought innovation to the field of ultrasonic monitoring. By integrating rigid ultras transducers with flexible substrates, newly developed ultrasonic devices not only exhibit stretchable and flexible characteristics but also conform closely to complex curved surfaces such as the human chest and joints [26, 27, 28, 29, 30, 31]. This design addresses the long-standing challenge of coupling traditional ultrasound probes with the skin while laying a solid foundation for the development of truly wearable ultrasound devices.

Significant progress has been made in this field of research. As early as 2018, Wang et al. pioneered a flexible ultrasound device that could conform tightly to the skin. This device was ultra-thin and stretchable, enabling non-invasive, continuous monitoring of blood pressure waveforms in human arteries and veins [32]. By 2024, an improved version of this device successfully underwent clinical validation on 21 patients, and the results confirmed the clear medical practicality and reliability of the flexible ultrasound device [33]. In 2021, Ding et al. developed a PMUT-based pulsed-wave Doppler ultrasound flowmeter, which monitored blood flow velocity changes via pulsed-wave Doppler mode and had a significantly smaller volume than traditional devices. However, it lacked flexible materials to conform to human skin effectively and featured complex PMUT manufacturing [34]. In 2025, Yang et al. further proposed a bandwidth-enhanced bone-shaped PMUT specifically designed for Doppler blood flow detection. This device utilizes a continuous-wave measurement mode, but its measurement accuracy somewhat depends on the precise fixation of the probe’s position. Therefore, in practical applications, operator correction is still required to ensure the reliability of the measurement results [35].

In recent years, the rapid advancement of flexible ultrasound technology has opened up new possibilities for non-invasive and continuous blood flow monitoring. Against this backdrop, this study proposes a wearable Doppler ultrasound patch based on flexible electronics technology, designed for real-time and continuous blood flow velocity monitoring. Specifically, by optimizing conventional microfabrication processes and incorporating serpentine electrode patterning techniques along with piezoelectric ceramic (PZT) composite material preparation, we have successfully developed a flexible ultrasound device with excellent stretchability and conformability. The patch integrates six ultrasound elements with a central frequency of 5 MHz, enabling tight adhesion to the human skin surface and accurately capturing various dynamic flow changes—from steady to pulsatile flows. This study aims to provide a high-accuracy, low-cost, and reliable solution for wearable Doppler blood flow monitoring devices, with potential applications in daily health management scenarios, thereby enhancing early screening and prevention capabilities for cardiovascular diseases.

## 2. Principle and structure

### 2.1 Pulsed-wave Doppler principle

The physical basis of ultrasonic Doppler blood flow measurement is the Doppler effect, which states that when there is relative motion between the wave source and the observer, the frequency of the wave received by the observer differs from the frequency emitted by the source. This frequency difference is known as the Doppler shift [36, 37]. The shift value contains critical information about blood flow velocity, and its quantitative relationship is described by the Doppler equation:

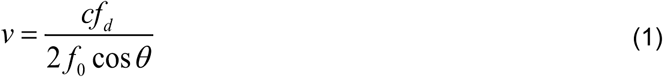

Among these variables,

*f_d_* represents the Doppler shift,

*f*_0_ denotes the center frequency of the ultrasound emitted by the transducer, *_v_* indicates the magnitude of the blood flow velocity vector, *θ* is the angle between the ultrasound beam and the direction of blood flow, and *c* represents the speed of ultrasound propagation in tissue. From this equation, it can be concluded that—provided the transmit frequency, speed of sound, and insonation angle are known—the absolute velocity of the target blood flow can be calculated by accurately measuring the Doppler shift.

Ultrasound Doppler technology is a core method for assessing hemodynamics, with pulsed-wave Doppler (PW Doppler) and continuous-wave Doppler (CW Doppler) representing two distinct operating modes based on different physical principles [38]. Their fundamental difference lies in how they handle the transmission and reception of sound waves. The physical principle of continuous-wave Doppler relies on the continuous emission and reception of sound waves. Its core mechanism lies in the fact that any Doppler-shifted signals—whether higher or lower than the transmitted frequency—generated by moving scatterers (such as red blood cells) along the beam path can be detected and processed simultaneously [39]. A direct consequence of this operating mode is that CW Doppler can measure extremely high blood flow velocities without an upper limit and does not suffer from signal aliasing, as the detection of frequency shifts is a continuous analog process. However, an inherent limitation of this principle is its lack of range resolution [40]. The system cannot distinguish the specific depth from which the frequency-shifted signals originate; all motion information along the acoustic beam path is superimposed and recorded as a composite signal, making it impossible to localize and analyze specific regions of interest.

In contrast, the physical principle of pulsed-wave Doppler is based on the pulse-echo and range gating techniques adopted from radar systems [41]. As shown in Fig. 1, the working principle of pulsed-wave Doppler receiving the original signal involves adjusting the opening time of the listening time window, allowing the operator to precisely select the depth of the sample volume, thereby enabling targeted blood flow assessment at specific anatomical locations. Pulsed-wave Doppler can determine the position of blood vessels based on the echo signal, which is more advantageous for subsequent signal processing and localization. Therefore, this study adopts this mode for measurement.

**Fig. 1.**
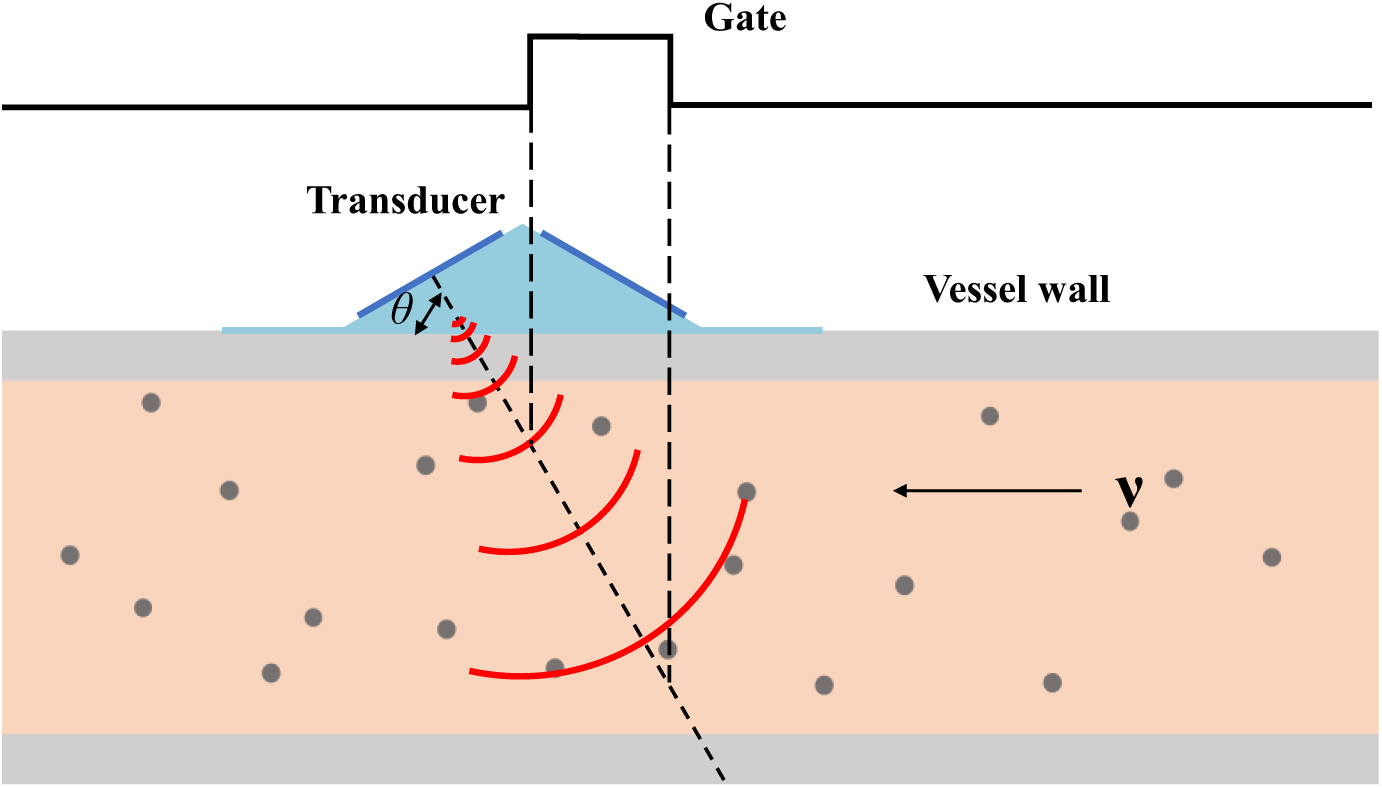
Pulsed-wave Doppler implements acquisition gating.

### 2.2 Ultrasound patch structure design

Fig. 2 illustrates the structural design of the developed wearable Doppler ultrasound patch. Its core feature lies in the use of a substrate with a specifically engineered tilt angle to precisely control the incident direction of ultrasonic waves. The patch integrates six ultrasonic transducers and two VIAs (Vertical Interconnect Access). These components are divided into two groups, each containing three transducers and one VIA, arranged symmetrically along the central axis. This layout ensures adequate coverage of the detection area by the patch while reducing reliance on precise manual alignment during measurement. Furthermore, by optimizing the number and arrangement of components, it effectively mitigates the cost increase and process complexity associated with using an excessive number of transducers. The device employs a meticulously layered top-down architecture: a polyimide (PI) film provides high stretchability, beneath which a copper (Cu) electrode layer interconnected with conductive adhesive (E-solder) forms a complete electrical pathway. A three-layer transducer design is adopted in this work. The piezoelectric ceramic serves as the core of the ultrasonic transducer, enabling bidirectional conversion between electrical and acoustic signals. The matching layer is acoustically optimized with an impedance value intermediate between the piezoelectric material and human tissue. The backing layer, attached behind the piezoelectric material, absorbs backward-propagating waves and dampens ringing effects. The most critical aspect of this design is the pre-defined angular substrate. Rather than being horizontally aligned, the substrate is precision-machined to a fixed inclination. This configuration dictates the transmission and reception direction of the integrated piezoelectric transducer array, establishing and maintaining a specific, non-perpendicular ultrasound incident angle (θ) relative to the skin surface. By locking this angle structurally, the design ensures that the ultrasonic beam consistently insonates the subcutaneous target vessels at an optimized and known θ, greatly reducing errors in blood flow velocity calculations caused by angle estimation uncertainty and significantly improving the accuracy and reliability of quantitative measurements.

**Fig. 2.**
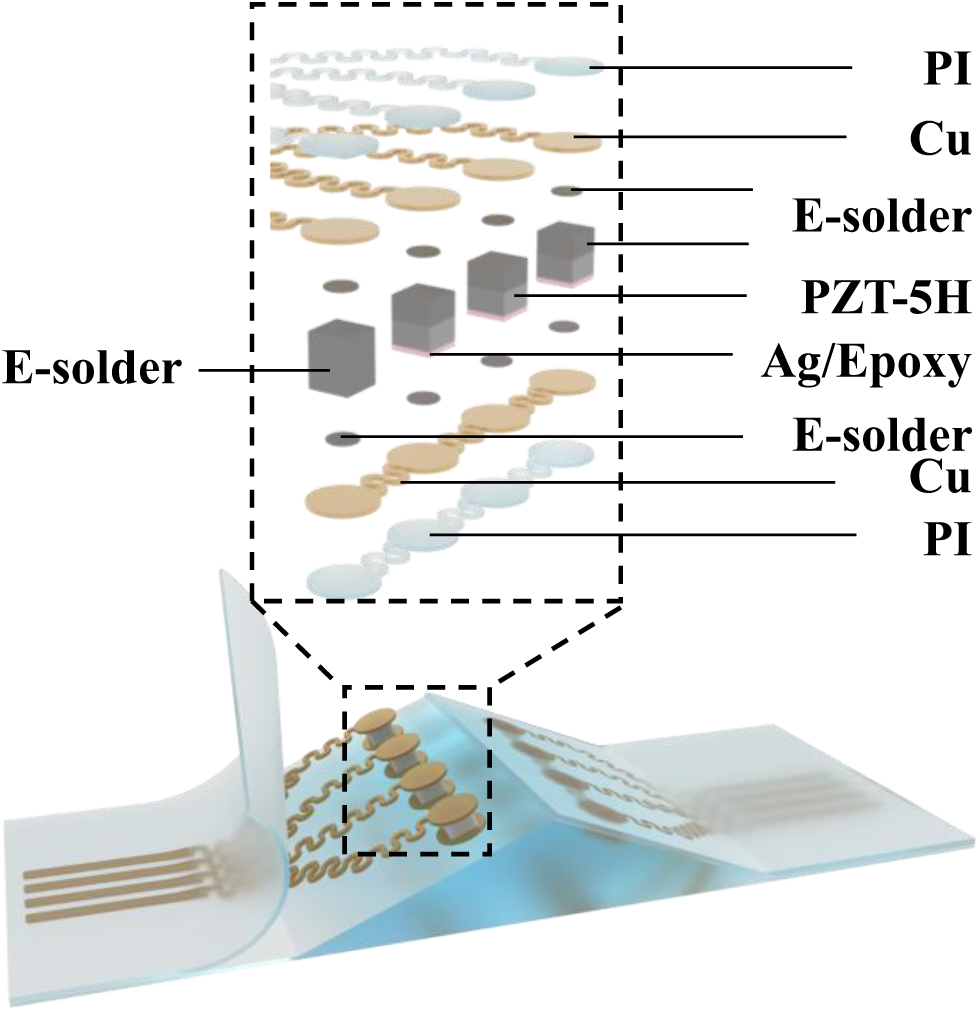
Structural diagram of the wearable Doppler ultrasound patch and exploded view of the ultrasonic transducers.

## 3. Materials and methods

### 3.1 Ultrasonic transducer design and fabrication

The three-layer ultrasonic transducer primarily consists of a matching layer, a piezoelectric layer, and a backing layer. In this study, Ag/Epoxy composite was selected for the matching layer, PZT-5H for the piezoelectric layer, and E-Solder 3022 for the backing layer. To optimize the performance of the fabricated transducer array, the Krimholtz, Leedom and Matthaei (KLM) equivalent circuit model was first employed for theoretical design and performance simulation. Based on one-dimensional transmission line theory, this model accurately describes the energy conversion behavior of the piezoelectric transducer at both electrical and acoustic ports. After balancing penetration depth and resolution, a center frequency of 5 MHz was chosen for the transducer. Using the KLM simulation platform, key parameters of each structural layer—including thickness, acoustic impedance, and acoustic attenuation—were systematically adjusted. The finalized parameters are summarized in Table 1.

**Table 1.**
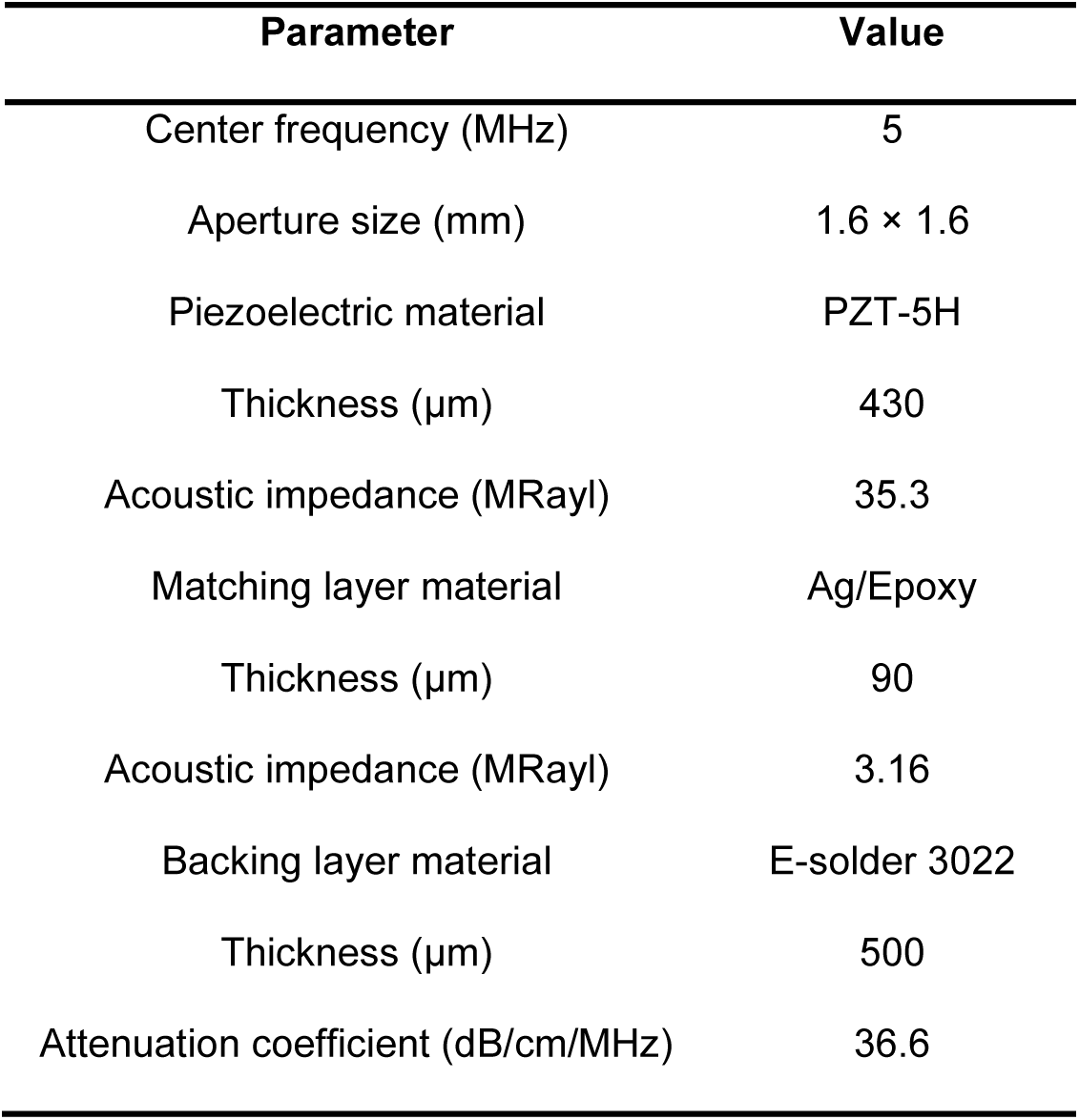
Parameters of the transducer elements.

The fabrication process of a single ultrasonic transducer is illustrated in Fig. 3. A PZT-5H piezoelectric ceramic measuring 1 cm × 1 cm with silver electrodes on both sides was selected as the base material. It was uniformly thinned to 430 μm via mechanical grinding. E-Solder 3022 (Von Roll USA Inc., Schenectady, NY, USA) was then applied to one side (with the original silver electrode retained) and cured at 45°C for 6 hours. The E-Solder 3022 layer was ground down to 500 μm to serve as the backing layer. A Cr/Au (500/1000 Å) electrode was deposited on the opposite side (without the retained electrode) using physical vapor deposition (Pro Line PVD 75, Kurt J. Lesker, USA). A matching layer was formed by mixing epoxy resin (Epo-Tek 301, Epoxy Technologies, Billerica, MA, USA) with high-purity silver particles 2–3 μm in diameter at a weight ratio of 1:2.4. This mixture was applied to the front surface of the piezoelectric layer, cured at 45°C for 6 hours, and then ground to a thickness of 90 μm. Finally, the transducer was diced into 1.6 mm × 1.6 mm units using a wafer dicing machine (DAD323, DISCO, Japan), completing the fabrication of the three-layer ultrasonic transducer.

**Fig. 3.**
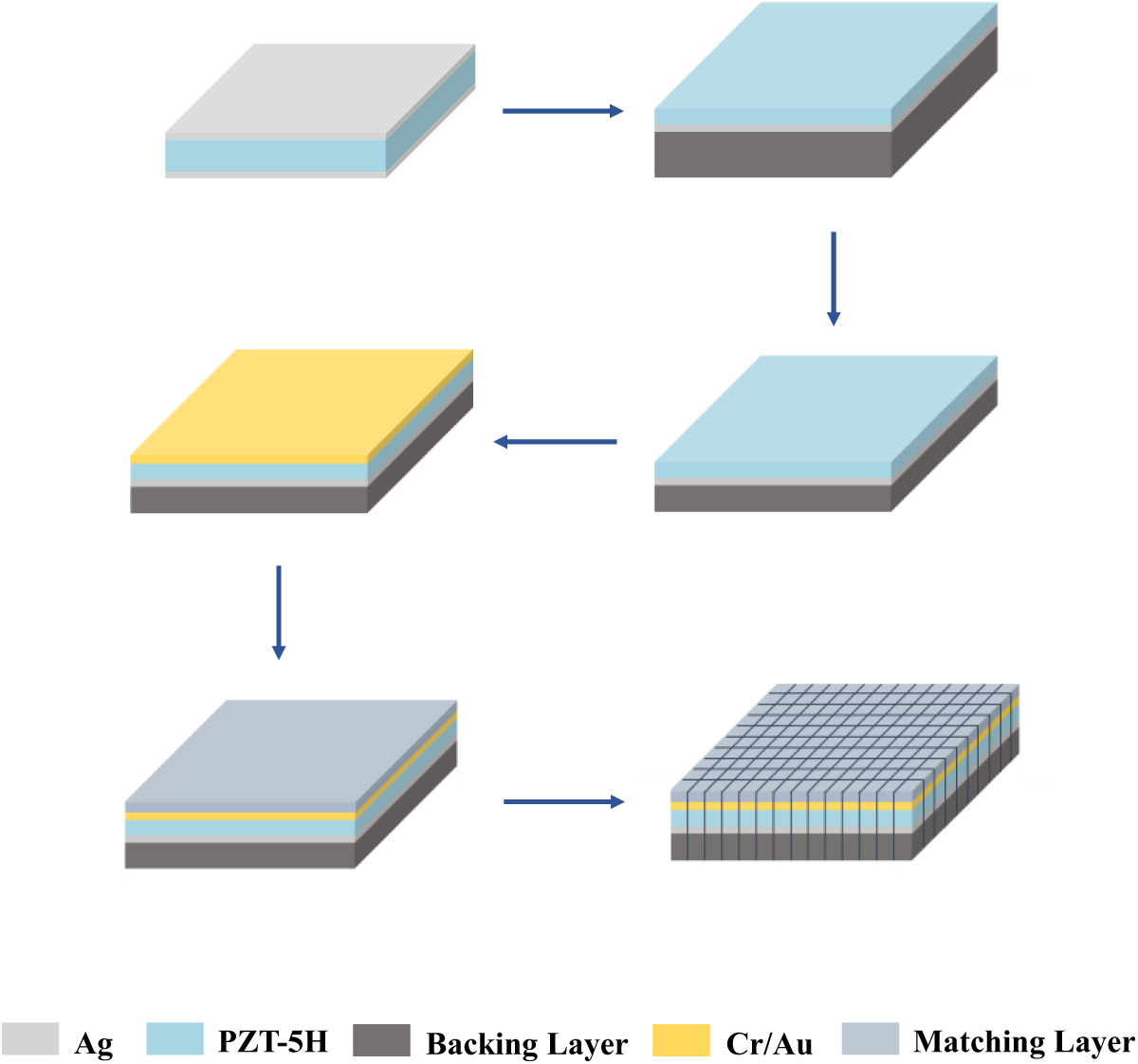
The fabrication process of a single ultrasonic transducer.

### 3.2 Electrode fabrication and flexible encapsulation

The serpentine electrode structure adopts a design with separated signal and ground traces, modeled using SolidWorks software (SolidWorks 2024, Dassault Systèmes, Waltham, MA, USA) with the specific configuration shown in Fig. 4. Fig. 4(a) shows the design dimensions of the serpentine electrode. The electrode features an "island-bridge" stretchable architecture. When subjected to bending or tensile deformation, strain is primarily concentrated on the flexible "bridge" structures, while the "island" regions remain largely undeformed, thereby protecting critical electrical connections from damage. The "bridge" structure is designed as an open-ring configuration with an arc opening angle of 120°, an outer diameter of 300 μm, and an inner diameter of 150 μm. This parameter set ensures compatibility with laser cutting precision requirements while providing the structure with favorable stretch recovery performance. The “island" region is designed as a complete circle with a radius of 0.8 mm, matching the dimensions of the ultrasonic transducer element. Due to the symmetrical arrangement of the transducers, the signal electrodes were designed in two mirrored versions, while the ground electrode employs a unified structure for general applicability. Fig. 4(b)-(d) show the designs of the signal electrode and grounding electrode. The trace width below the signal electrodes is 0.5 mm, with an inter-line spacing of 1 mm, matching the 1 mm pitch FFC (flexible flat cable) for easy integration and connection.

**Fig. 4.**
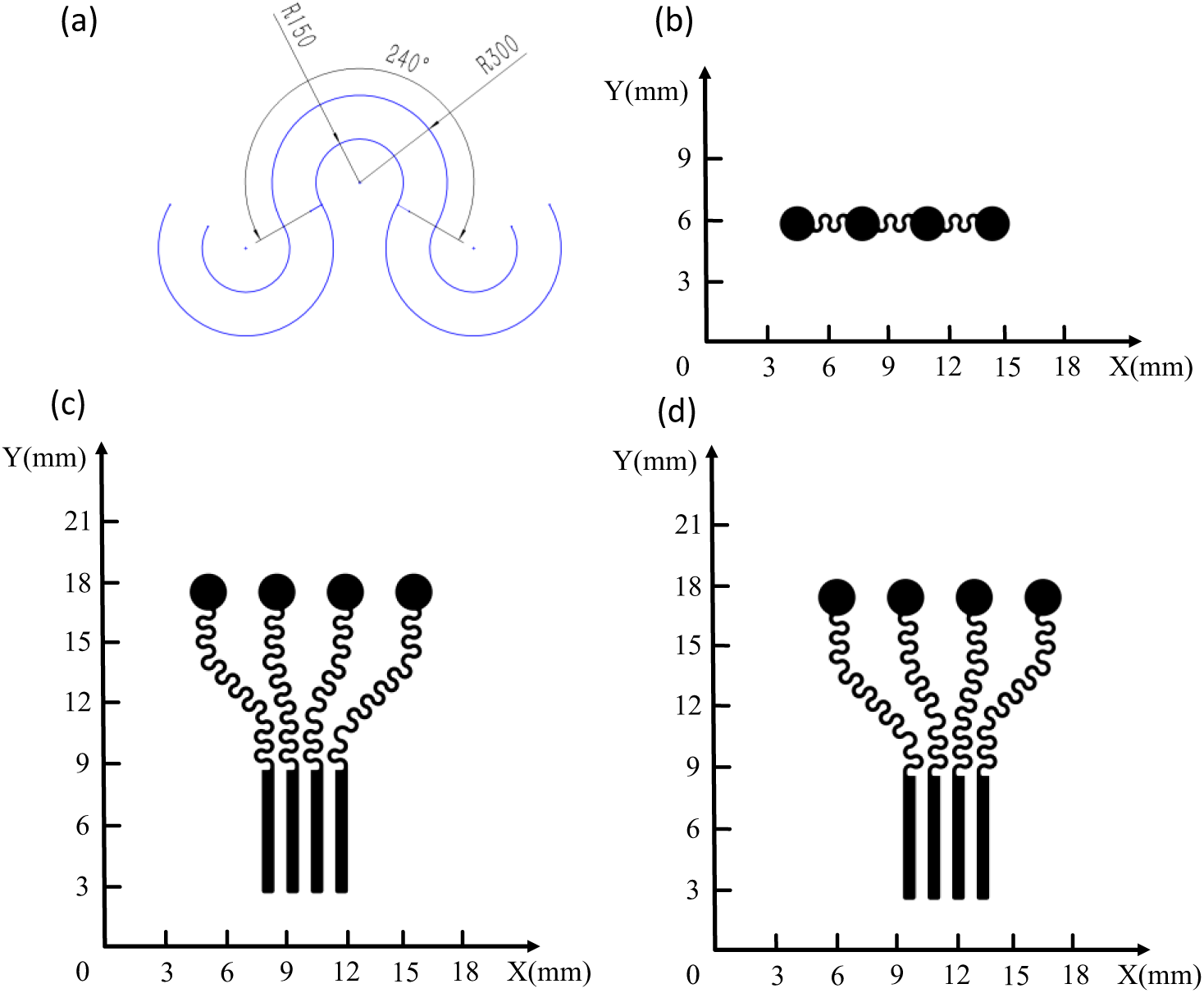
(a) Serpentine electrode design dimensions, (b) grounding electrode design, (c) signal electrode design, and (d) symmetric signal electrode design.

The process steps for electrode processing and flexible packaging are shown in Fig. 5, where our technique avoids the transfer printing and spin-coating of Cu and PI. A PVC board was first selected as the substrate. Ecoflex was spin-coated (1000 rpm, 1 minute) onto it and cured at room temperature for 2 hours. After curing, a copper foil (Cu: 10 μm, PI: 12 μm) was placed and firmly laminated onto the surface. A laser cutter was then used to directly pattern the foil without a mask. After patterning, an anisotropic conductive adhesive was applied to connect the FFC to the signal electrodes. E-Solder 3022 was used to connect the backing layer side of the transducers, followed by curing in an oven at 45°C for 6 hours. Subsequently, E-Solder 3022 was again used to connect the matching layer side of the transducers to the ground electrode. The signal electrode contains four "islands," with the fourth island connected to a VIA. This VIA was formed by thinning cured E-Solder 3022, serving to bring the ground electrode to the same plane as the signal electrode for routing through the FFC. Finally, Ecoflex was poured to encapsulate the entire assembly. After curing at 45°C for 1 hour, the PVC substrate was removed, yielding two symmetric flexible patches.

**Fig. 5.**
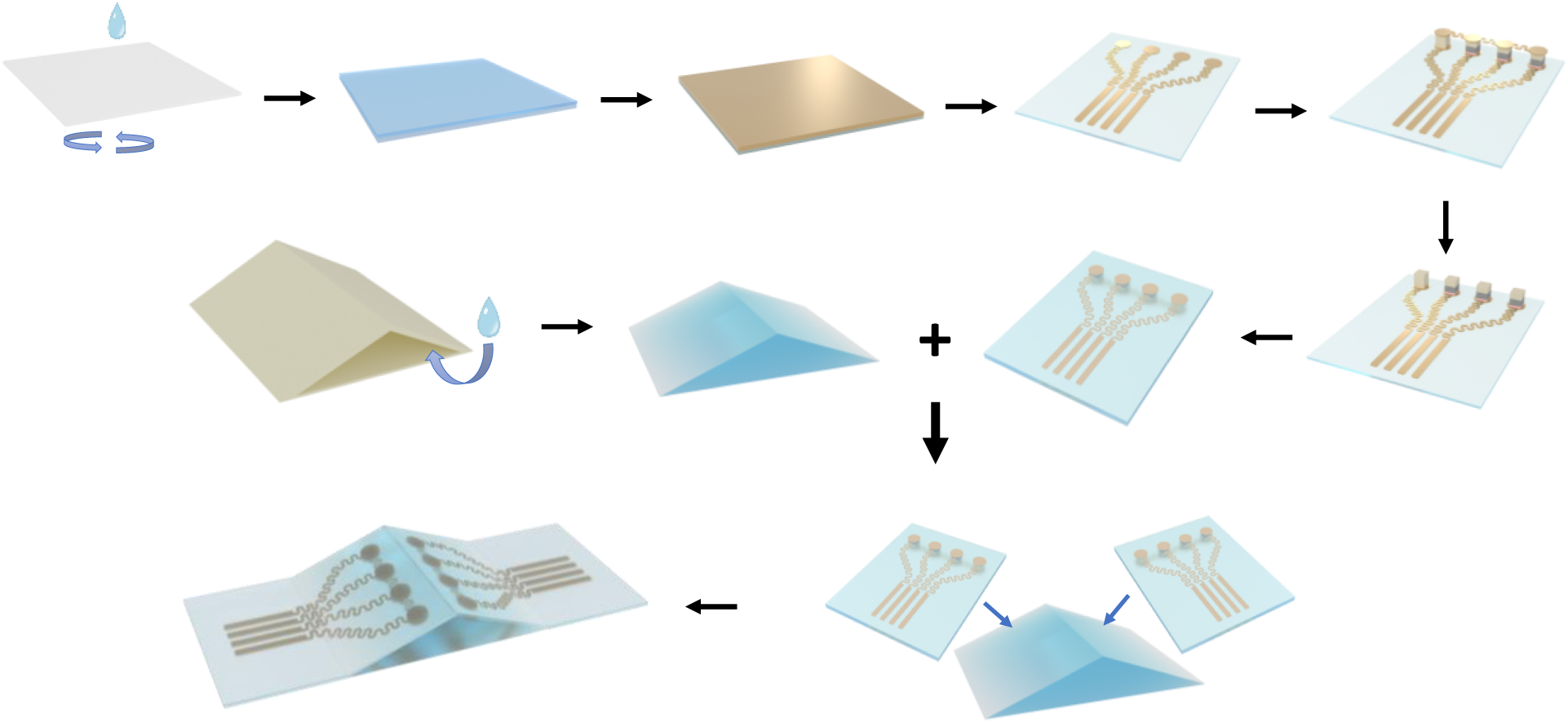
The complete process of serpentine electrode patterning and flexible encapsulation in the manufacturing of ultrasound patch.

A mold with a specific structure was fabricated using 3D printing. Ecoflex silicone was poured into the mold cavity, followed by vacuum degassing to eliminate air bubbles and ensure a defect-free internal structure. After curing, the mold was removed to obtain a flexible substrate with a 30° tilt angle. The pre-prepared wearable ultrasound patch was then symmetrically positioned and fixed onto this substrate. Ecoflex was poured again to fully encapsulate the entire structure and cured at 45°C for 1 hour to achieve final integration. Through the above process, we successfully fabricated a wearable ultrasound patch device with a predetermined tilt angle.

### 3.3 Performance measurements

Upon completion of the wearable ultrasound patch fabrication, we systematically evaluated its electrical performance, acoustic characteristics, and mechanical tensile properties. Specific experiments included: using an impedance analyzer (E4990A, Keysight Technologies, USA) to measure the resonant frequency and impedance amplitude of the ultrasonic transducers in the patch; and performing pulse-echo experiments to determine the center frequency and -6 dB bandwidth. All tests were conducted under the initial non-stretched state of the patch to establish a performance baseline. Additionally, the stretchability of the patch was quantitatively characterized through uniaxial tensile tests.

In the pulse-echo response test, the wearable ultrasound patch was fixed onto a plastic substrate and fully immersed in degassed water. A steel block was placed directly in front of it as a reflective target to generate measurable echo signals. The experiment utilized a commercial pulser-receiver (DPR 500, JSR Ultrasonics, NY, USA) for excitation and reception. The echo signal was amplified with a 30 dB gain and acquired by a high-bandwidth commercial oscilloscope (DSOX3054G, Keysight Technologies, USA). The built-in spectrum analysis function of the oscilloscope was further employed to transform the echo signal into the frequency domain, from which the spectrum was obtained and key parameters such as the -6 dB bandwidth and center frequency were calculated. For mechanical characterization, uniaxial tensile tests were conducted to evaluate the patch’s stretchability. Axial tension was gradually applied to the patch while its deformation process was observed and the maximum stretch length was recorded.

To further validate the capability of the developed transducer to capture blood flow velocity, an in vitro experiment was conducted, with the experimental setup shown in Fig. 6. The blood flow phantom consisted of a silicone tube with an inner diameter of 5.5 mm, through which Doppler fluid (Model 769DF, CIRS, USA) was circulated using a flow pump (LKTC-L CONTROLLER, Preclinic, China) to simulate realistic blood flow. Excitation signals were generated by the DPR 500, and the echo signals were acquired by a data acquisition system (PXIe-1092, NI, USA). The acquisition channel of the patch was controlled via an electronic switch. The experimental parameters were set as follows: pulse repetition frequency (PRF) of 10 kHz, sampling frequency of 250 MHz, and a fixed Doppler incident angle of 60°. To systematically evaluate the performance of the patch, the experiment was divided into two parts: first, testing the accuracy of the patch under different constant flow velocities; second, validating its ability to capture dynamically varying flow rates through pulsatile flow tests. Control groups with corresponding flow rates were established for all experiments to ensure reliability and comparability of the results.

**Fig. 6.**
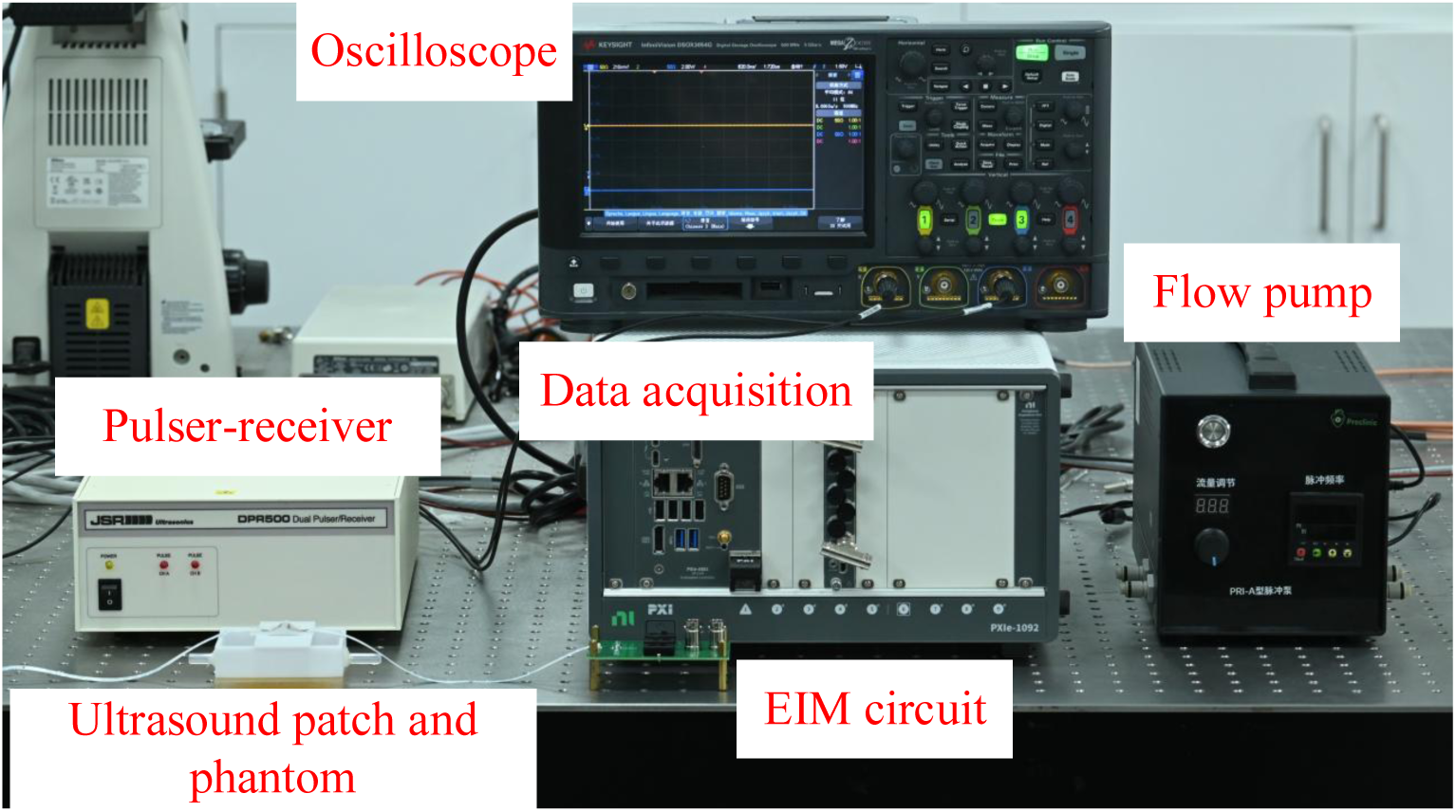
Description of the in vitro experimental setup.

### 3.4 Electrical impedance matching

An ultrasonic transducer can be modeled as an equivalent circuit consisting of resistors, capacitors, and inductors. Its equivalent circuit near the resonant frequency is shown in the Fig. 7(a). In the circuit, C₀ represents the static capacitance between the two electrodes, Cₘ is the dynamic capacitance, Lₘ is the dynamic inductance, and Rₜ is the dynamic resistance. The dynamic resistance Rₜ comprises two components: Rₘ, which represents the mechanical loss of the transducer, and Rₗ, which denotes the radiation load resistance accounting for energy radiated into the medium. The admittance formula of this equivalent circuit is given by:

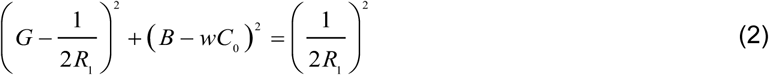

**Fig. 7.**
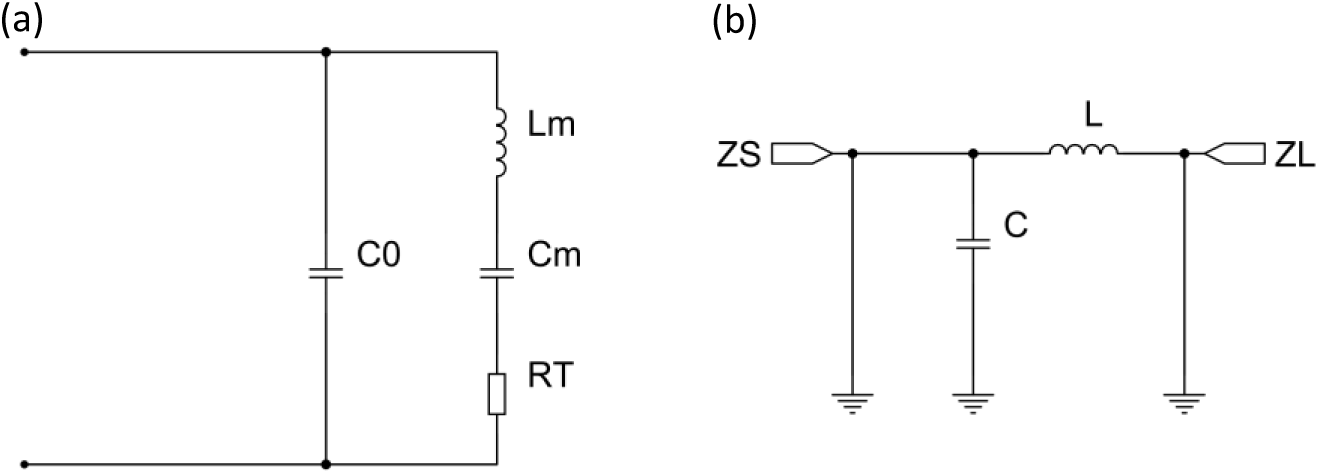
(a) Equivalent circuit of ultrasonic transducer, and (b) L-type matching network structure.

In the equation, *G* represents the equivalent conductance and *B* represents the equivalent susceptance [42].

At the series resonant frequency (fₛ), the equivalent susceptance of the transducer becomes zero, and its impedance appears purely resistive. This resistance value corresponds to the dynamic resistance (Rₜ) of the transducer. If the dynamic resistance (Rₜ) matches the internal resistance (Rₛ) of the signal source (Rₜ = Rₛ), the condition for the maximum power transfer theorem is satisfied. This means that the power delivered by the source can be most efficiently absorbed by the transducer and converted into mechanical vibration or acoustic energy. After characterizing the impedance properties and resonant frequency of the self-fabricated ultrasonic transducer using an impedance analyzer, an L-type matching network structure, as shown in Fig. 7(b), was employed for impedance matching. The designed transducer has a center frequency of 5 MHz and requires impedance matching with a DPR500 pulser–receiver, which has an internal resistance of 50 Ω. By calculating the required capacitance and inductance parameters for the matching network, a dedicated PCB was designed and fabricated. The transducer was then connected to this circuit to achieve impedance matching.

### 3.5 Slow-time sampling theorem

In ultrasonic Doppler blood flow measurement, quadrature demodulation is a classical technique wherein two reference signals with a 90° phase difference are generated to downconvert the echo signal through mixing and low-pass filtering, ultimately extracting a complex baseband signal (I and Q channels) that retains complete amplitude and phase information. Although this method effectively preserves the spectral characteristics of blood flow signals, its high computational complexity and intricate processing pipeline render it inefficient—particularly in large-scale data scenarios or applications demanding real-time performance. To overcome these limitations, we introduced a more efficient processing strategy: slow-time sampling [43].

The specific algorithm flow is shown in Fig. 8: First, the raw one-dimensional acquired signal undergoes pulse separation and temporal alignment to extract individual complete pulse-echo segments. A bandpass filter is then applied to suppress irrelevant frequency components and noise, preserving the effective Doppler signals generated by red blood cell scattering. Next, a specific depth location is selected, and the amplitude at this location within each pulse echo is sequentially extracted to form a slow-time-varying signal sequence. This slow-time signal is then processed using a Hilbert transform to extract its envelope, followed by Fast Fourier Transform (FFT) to analyze its spectral distribution. Prominent peaks in the spectrum correspond to the dominant flow velocity of the blood. To construct a two-dimensional spatiotemporal distribution map, the algorithm iterates the above steps across all depth points, integrating frequency components at various time instants and depths to form a two-dimensional frequency-time matrix. This matrix undergoes dynamic threshold binarization to emphasize valid Doppler components, followed by mean filtering to suppress discrete noise. Finally, spline interpolation is applied to smooth and optimize the binarized result, yielding a clear and accurate binary image of blood flow velocity distribution that provides a reliable foundation for subsequent quantitative analysis.

**Fig. 8.**
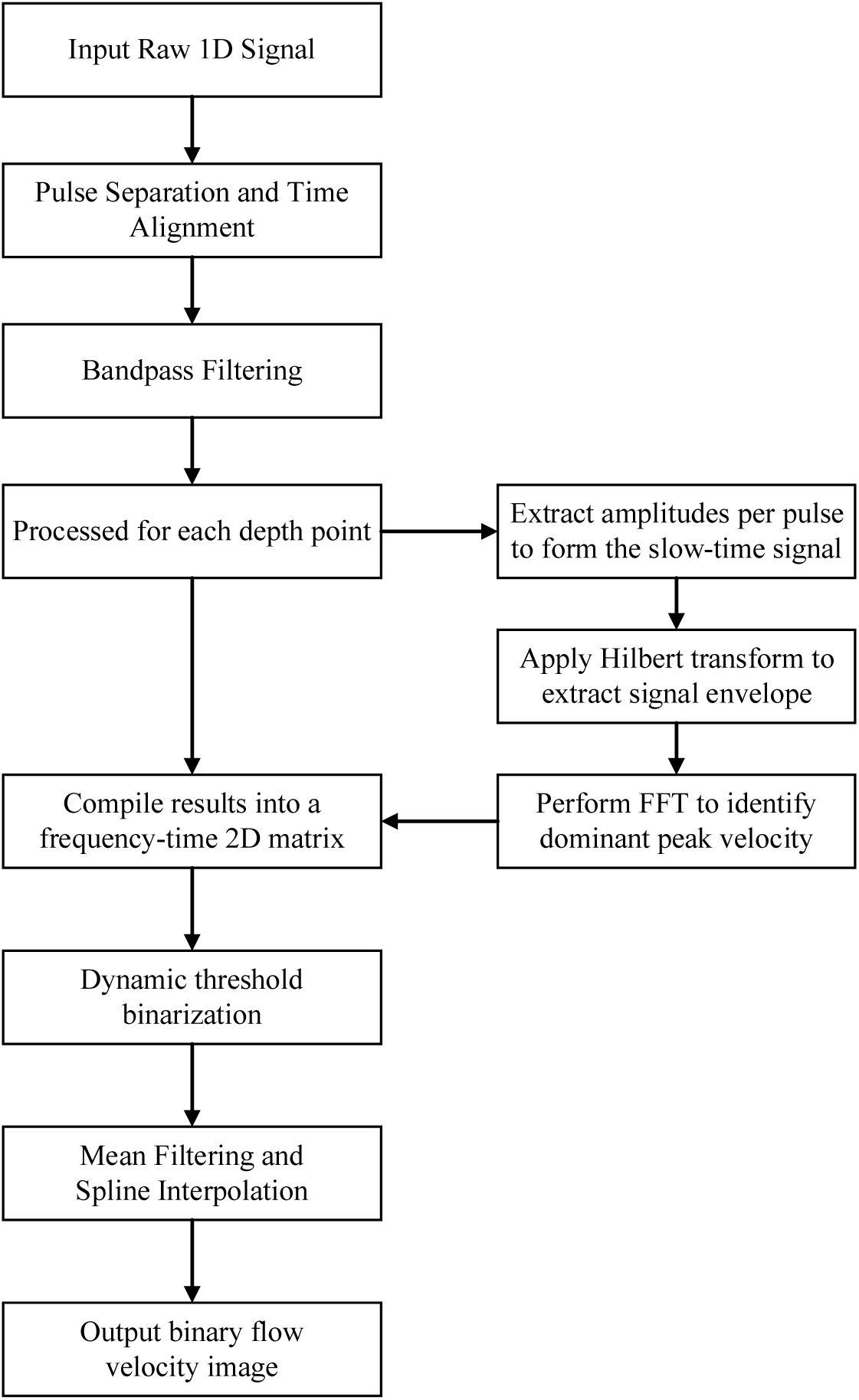
Slow-time sampling algorithm processing flow.

## 4. Results and discussion

### 4.1 Doppler ultrasound patch prototype

Fig. 9(a) displays the physical prototype of our developed wearable Doppler ultrasound patch. The patch features a 2 cm × 4 cm core area, and a weight of 2.01 g. Two sets of transducers are routed via separate FFC (Flexible Flat Cable) connections, and the entire assembly is encapsulated in Ecoflex material, ensuring flexibility and durability. Fig. 9(c)-(d) demonstrate the patch’s conformability when attached to curved surfaces with different radii. Fig. 9 (b) illustrates the patch’s ability to adhere closely to the human skin surface without requiring additional hydrogel. Thanks to its integrated serpentine electrode structure, the patch can stretch and extend synchronously with skin motion, effectively maintaining signal transmission stability. The substrate, precision-machined with a 30° tilt angle, ensures that the patch maintains a Doppler incident angle of approximately 60° even in bent states, thereby guaranteeing accurate blood flow velocity measurement. Furthermore, the inter-transducer spacing within the patch has been optimized to 3.2 mm, a configuration designed to maximize acoustic coverage of the target blood vessels.

**Fig. 9.**
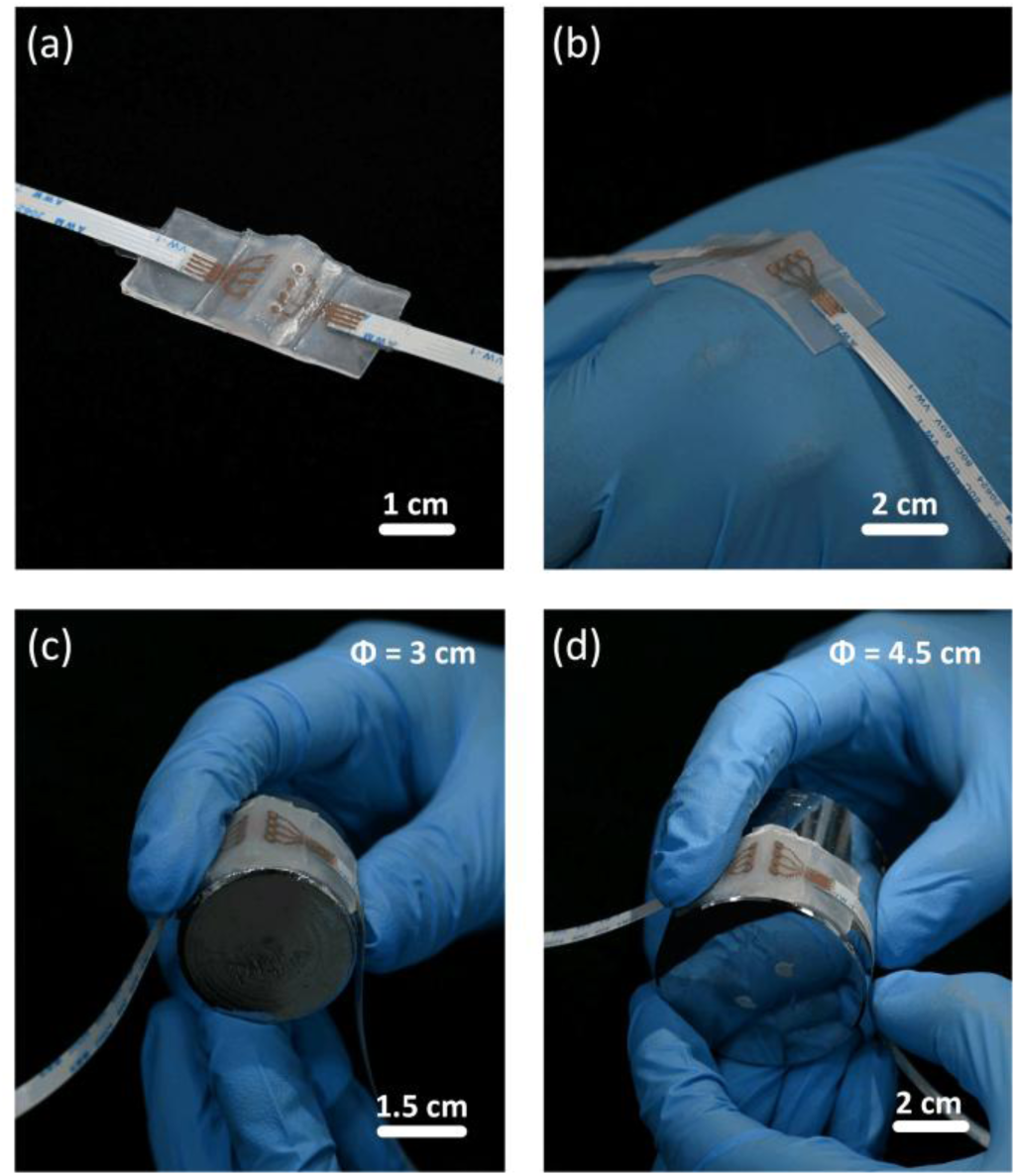
Wearable Doppler ultrasound patch (a) physical display, (b) closely attached to human skin, (c) attached to a metal cylinder with a diameter of 3 cm, and (d) attached to a metal cylinder with a diameter of 4.5 cm.

### 4.2 Basic performance of the ultrasound patch

Based on the KLM simulation model, the optimization design results are shown in Fig. 10. Fig. 10(a) shows the impedance spectrum of the transducer obtained through simulation, and Fig. 10(b) shows the corresponding pulse-echo impulse response. The simulation results indicate that the designed ultrasonic transducer has a center frequency of 5.0 MHz, a -6 dB bandwidth of 46.5%, and a resonant frequency of 4.85 MHz.

**Fig. 10.**
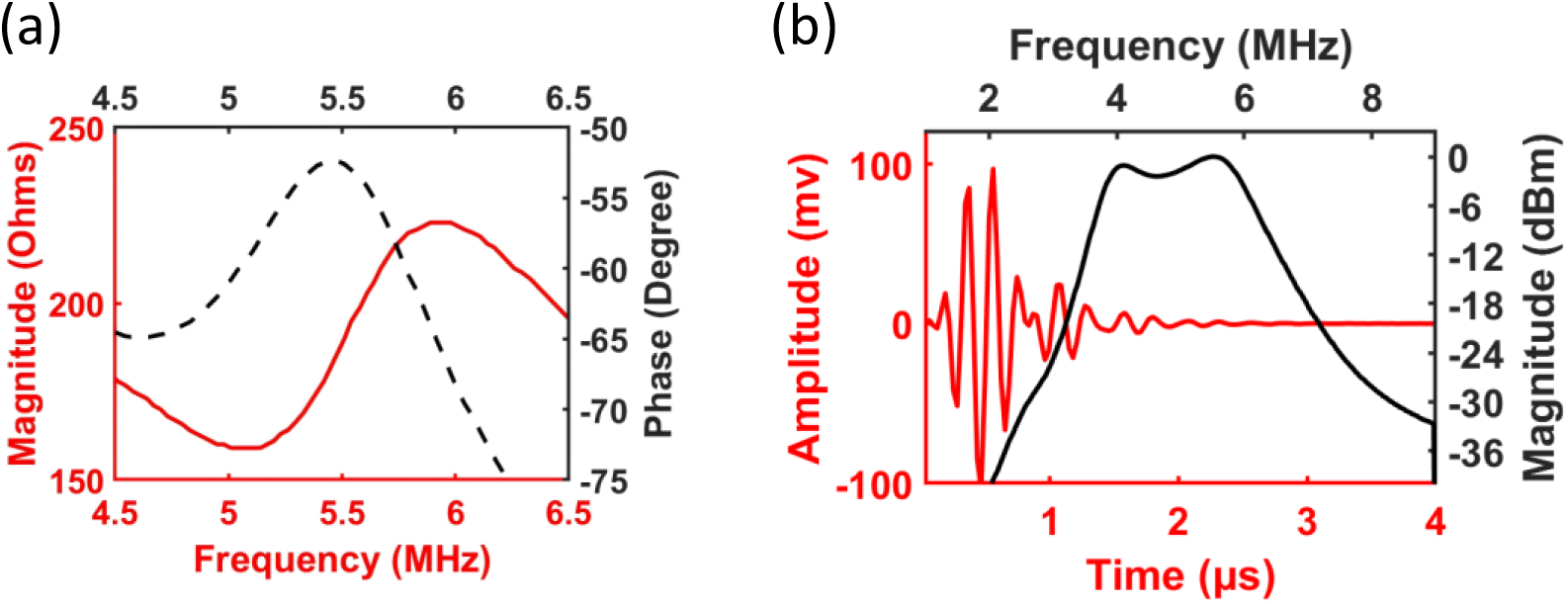
Under KLM simulation: (a) impedance spectrum, and (b) pulse-echo response.

After completing the patch fabrication, impedance matching and performance testing were conducted. The measured impedance spectra of the six transducers after impedance matching are shown in Fig. 11(a), and the pulse-echo impulse response of one representative transducer is shown in Fig. 11(b). The test data show that this representative transducer has a center frequency of 5.05 MHz, a -6 dB bandwidth of 42.58%, an impedance spectrum indicating a resonant frequency of 4.9 MHz, and an impedance value of 165 Ω after matching.

**Fig. 11.**
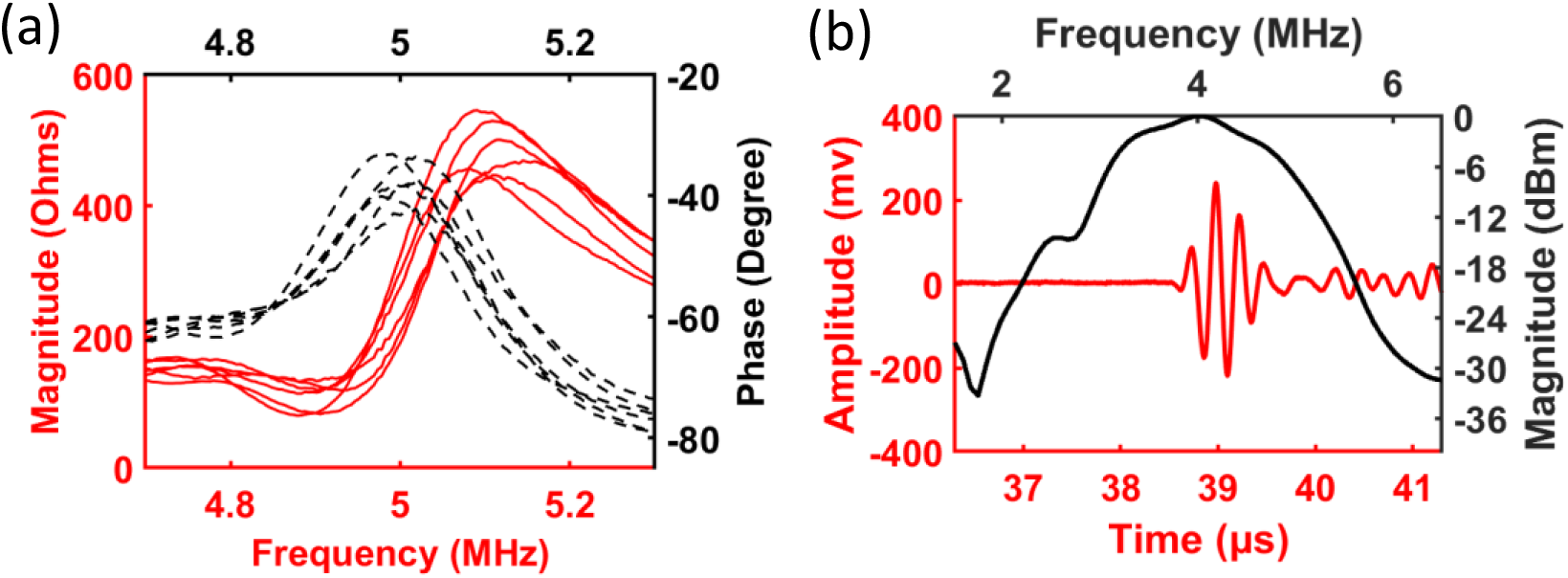
Experimental measurements of (a) impedance spectrum, and (b) pulse-echo response.

Performance testing was conducted on all ultrasonic transducers, and the obtained key parameters, including center frequency, -6 dB bandwidth, resonant frequency, and impedance value, are shown in Fig. 12. Specifically, Fig. 12(a) shows that the center frequency distribution range of the transducers is 5.05 ± 0.04 MHz; Fig. 12(b) shows the corresponding -6 dB bandwidth range of 44.06 ± 3.07%; Fig. 12(c) presents the resonant frequency range after impedance matching as 4.92 ± 0.04 MHz; and Fig. 12(d) gives the impedance value range of the transducers after matching as 131 ± 34 Ω. Changes in the impedance values of each ultrasonic transducer after impedance matching can be attributed to several factors. First, discrepancies between theoretical calculations and actual measurements prevent ideal matching outcomes, with PCB parasitic capacitance further influencing the results. Second, inherent variations in transducer manufacturing and assembly reduce overall consistency.

**Fig. 12.**
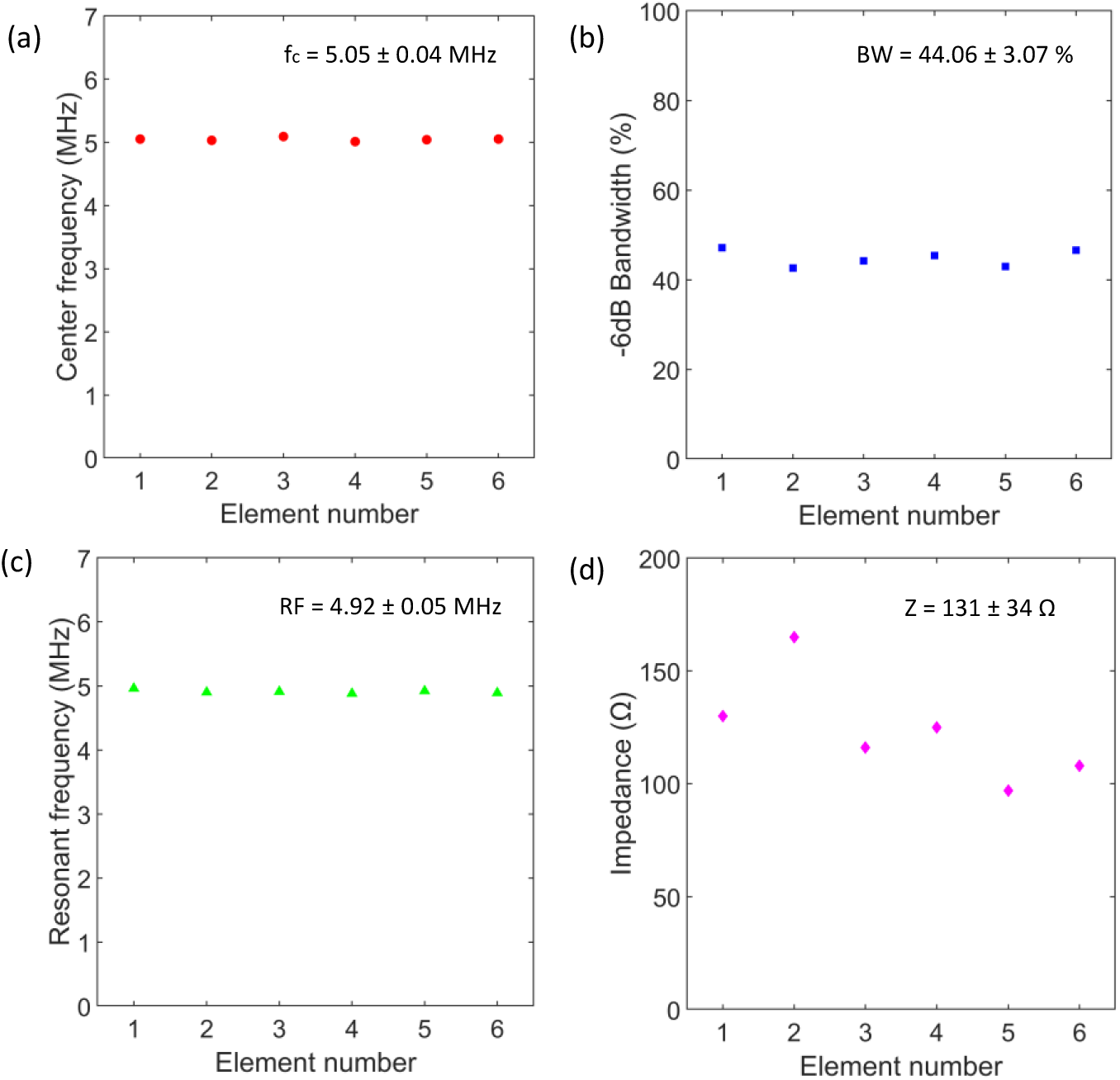
The experimental measurements of the ultrasound patch are translated as follows: (a) center frequency, (b) -6 dB bandwidth, (c) resonant frequency, and (d) impedance value.

A uniaxial tensile test was performed on the ultrasonic patch (results shown in Fig. 13). The experiment results show that the patch can achieve a maximum elongation of 30%. Benefiting from this mechanical characteristic, the device can form a stable and seamless contact with human skin. This is because human skin typically exhibits a linear elastic response when tensile strain is less than 20%, and the patch’s extensibility fully adapts to this mechanical characteristic [44].

**Fig. 13.**
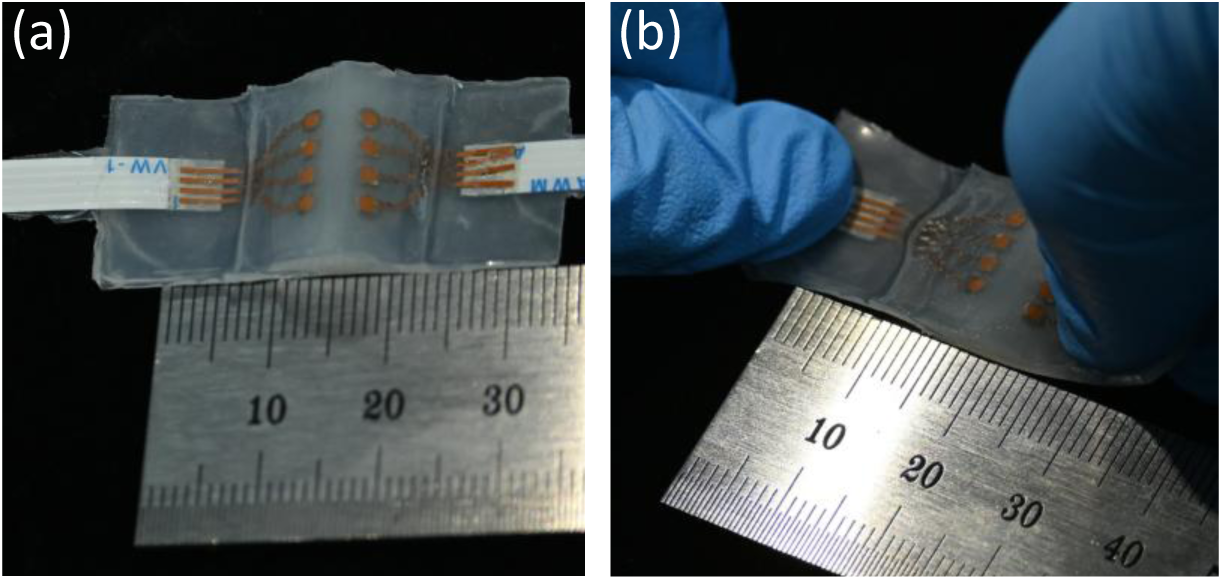
Ultrasound patch under (a) unstrained state and (b) 30% stretched condition.

### 4.3 In vitro flow velocity measurement

An in vitro experiment was conducted to validate the blood flow measurement capability of the transducer. Fig. 14 displays the echo signal received by a typical transducer element. The front and back walls of the phantom tube are clearly distinguishable based on the echo amplitude, with the target fluid region located between them. This provides crucial target reference information. During the experiment, the echo signal was monitored in real-time using an oscilloscope. The optimal transducer channel was selected by switching channels, followed by data acquisition. The acquired data underwent slow-time sampling processing to obtain the flow velocity measurement results.

**Fig. 14.**
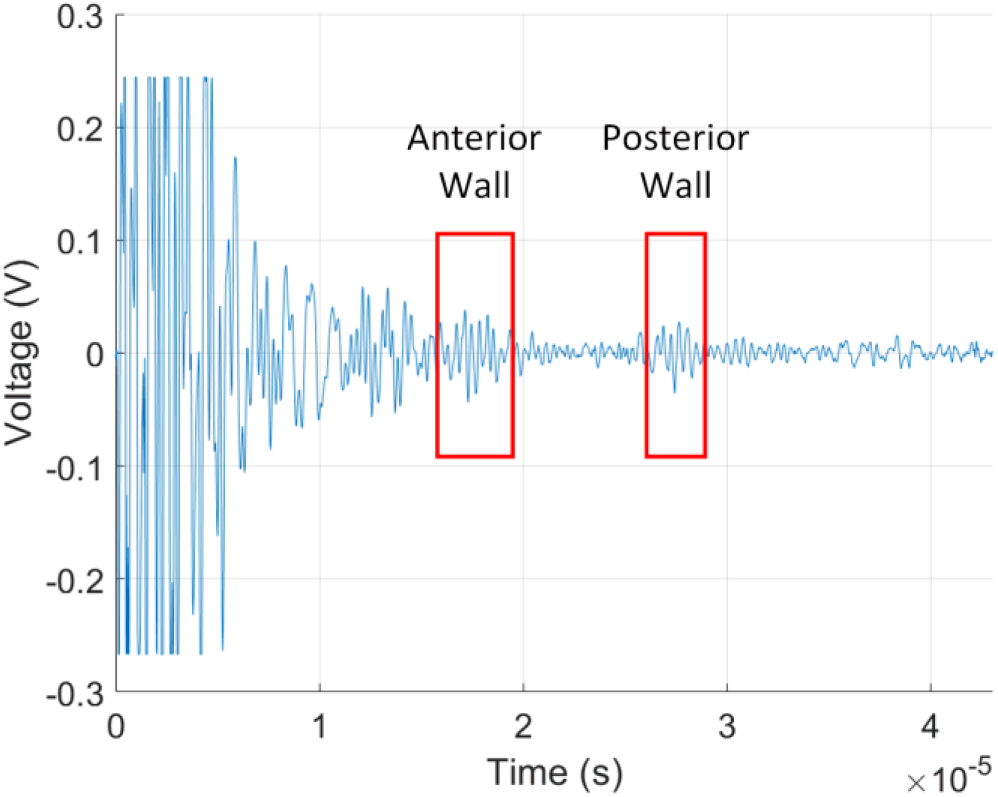
The echo signal received by a typical transducer element.

We first conducted constant-speed measurement experiments, with the flow velocity standard defined by the parameters set on the flow pump. Fig. 15(a) to (c) display the results for constant flow velocities of 50 cm/s, 60 cm/s, and 70 cm/s, respectively. Subsequently, pulsatile flow velocity measurement experiments were performed. Due to limitations in the flow pump’s performance, experimental groups were designed with pulsation frequencies of 1 cycle/s and 2 cycles/s, each corresponding to peak flow velocities of 50, 60, and 70 cm/s. The results are shown in Fig. 15(d) to (f): Fig. 15(d) represents a peak velocity of 50 cm/s with a pulsation of 1 cycle/s, Fig. 15(e) corresponds to 60 cm/s at 2 cycles/s, and Fig. 15(f) to 70 cm/s at 1 cycle/s.

**Fig. 15.**
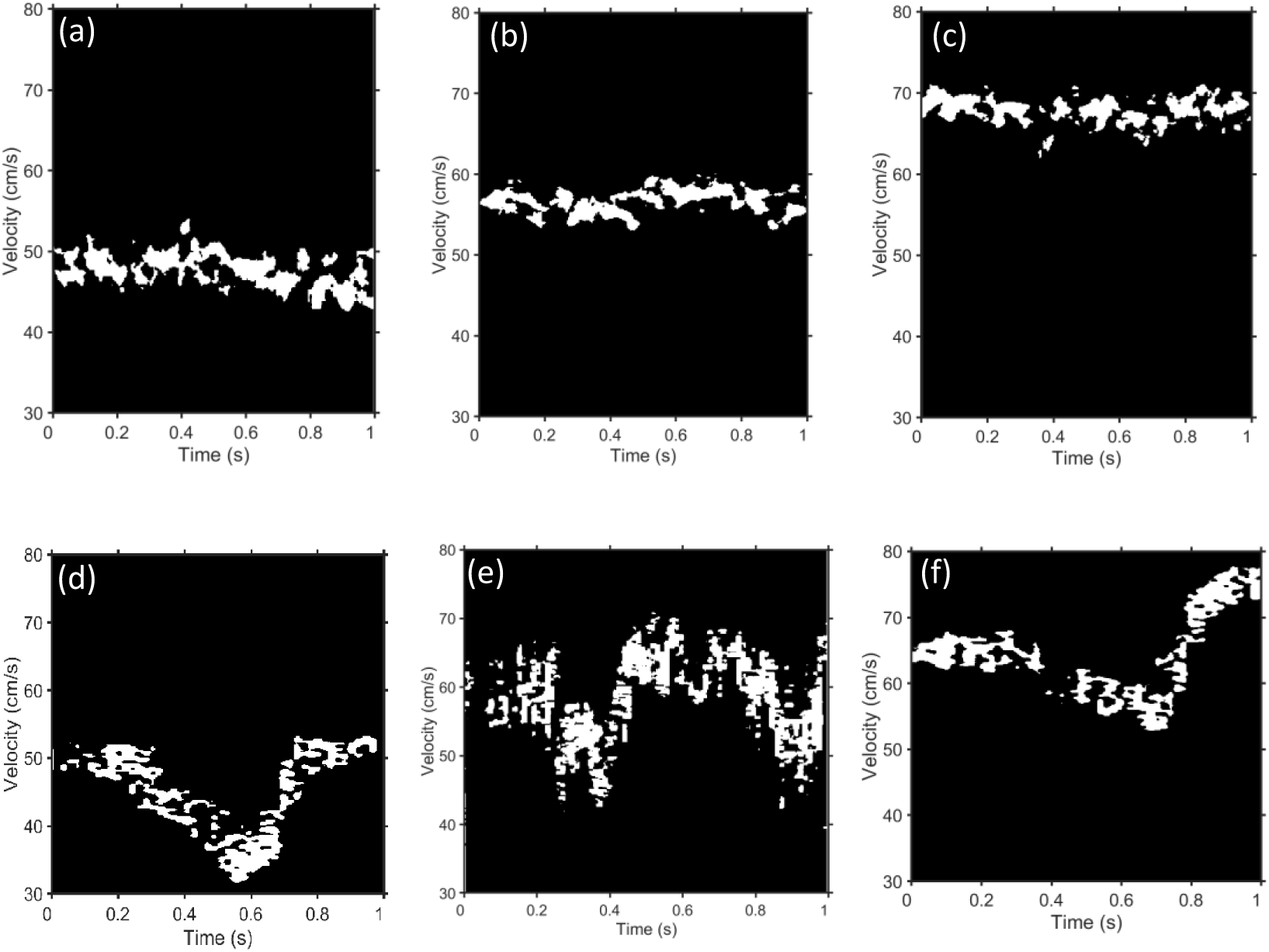
In the in vitro flow velocity experiment, constant velocities: (a) 50 cm/s, (b) 60 cm/s, (c) 70 cm/s, and pulsatile velocities: (d) a peak velocity of 50 cm/s with a pulsation of 1 cycle/s, (e) a peak velocity of 60 cm/s at 2 cycles/s, and (f) a peak velocity of 70 cm/s at 1 cycle/s.

To further assess the measurement accuracy of the ultrasound patch, a quantitative analysis was applied to the constant-speed experimental group. The specific method involved extracting the maximum and minimum values from the flow velocity regions marked in the figures, calculating their average, and comparing it with the reference value set by the flow pump to analyze the systematic error. The results are summarized in Table 2. The absolute percentage error ranges from 4.1% to 12%, and the true flow velocity ranges from 20 to 100 cm/s. The mean absolute error (MAE) is 3.56 cm/s, and the mean absolute percentage error (MAPE) is 5.99%. Table 3 shows the measured blood flow velocity values in the carotid artery of adult males and females. As blood flow in human arteries is dynamically changing, the measured velocities fluctuate correspondingly across different cardiac phases [45]. Here, d represents the end-diastolic velocity, S1 denotes the peak systolic velocity, S2 indicates the second systolic velocity, I corresponds to the incisura between systole and diastole, and D signifies the peak diastolic velocity. The measured blood flow velocities in this experiment range from 20 cm/s to 100 cm/s, covering the main velocity range observed in the carotid artery [46]. This result further supports the promising potential of the wearable ultrasound patch developed in this study for hemodynamic monitoring applications.

**Table 2.** Comparison of measured velocity and actual velocity.

| Actual Velocity (cm/s) | Measured Velocity<br>(cm/s, Mean ±SD) | Absolute percentage<br>error (% , Mean) |
| --- | --- | --- |
| 20 | 17.6±2.7 | 12 |
| 30 | 26.9±3.0 | 10.33 |
| 40 | 36.9±3.7 | 7.75 |
| 50 | 47.1±3.6 | 5.8 |
| 60 | 57.4±3.4 | 4.33 |
| 70 | 67.1±3.8 | 4.14 |
| 80 | 76.1±3.7 | 4.88 |
| 90 | 85.1±4.1 | 5.44 |
| 100 | 94.9± 4.8 | 5.1 |

**Table 3.** Flow velocity and data in women and men.

| Flow velocity data | Women | Men |
| --- | --- | --- |
| d (cm/s) | 23.7±0.8 | 20.9±0.7 |
| S1 (cm/s) | 96.9±3.2 | 91.2±3.1 |
| S2 (cm/s) | 64.3±2.0 | 50.0±1.8 |
| I (cm/s) | 35.6±1.3 | 27.5±0.9 |
| D (cm/s) | 45.6±1.2 | 39.1±1.0 |

However, the experimental results still exhibit certain measurement errors. Firstly, influenced by the viscous effects of the vessel wall, the velocity profile of blood flow within the vessel is parabolic, with higher velocities at the center and lower velocities near the wall. This non-uniform velocity distribution leads to an inevitable discrepancy between the measured values and theoretical expectations. Secondly, the non-uniformity in the concentration, size, and spatial distribution of the scattering particles within the phantom, which are used to reflect the ultrasonic signals, causes fluctuations in the Doppler frequency shift signal, thereby affecting the accuracy of the flow velocity inversion results. Furthermore, the presence of strong electromagnetic interference in the experimental environment introduces additional noise, which not only causes signal distortion and data fluctuations but also represents a major limiting factor for transitioning the current system to human studies.

## 5. Conclusion

This study successfully developed a wearable Doppler ultrasound patch for continuous blood flow monitoring. The patch is entirely encapsulated with flexible materials, with a core area of only 2 cm × 4 cm and a weight of 2.01 g, effectively overcoming the limitations of traditional probes being rigid and bulky. In terms of hardware design, the patch integrates six ultrasonic transducers and two E-Solder posts; each transducer consists of a matching layer, a piezoelectric layer, and a backing layer, with a center frequency of 5.05±0.04 MHz, a -6 dB bandwidth of 44.06±3.07%, a resonant frequency after impedance matching of 4.92±0.04 MHz, and an impedance value of 131±34 Ω.

The core innovations of this design include two aspects: first, simplifying the traditional serpentine electrode processing technique, avoiding the complex transfer printing and spin-coating of Cu/PI steps; second, adopting a substrate structure with a preset angle, fixing the tilt angle through precision machining, locking the specific incident angle (θ) of the ultrasound wave relative to the skin surface, thereby ensuring that the acoustic beam incidents into the subcutaneous target blood vessels at an optimal and known angle, significantly improving the accuracy and reproducibility of measurements. In terms of signal processing, the study employs slow-time sampling for blood flow frequency shift extraction, which significantly reduces computational costs while maintaining accuracy. Through in vitro experiments, it was verified that the patch can accurately measure constant-speed fluid velocity and effectively capture the dynamic changes of pulsatile flow, demonstrating its feasibility in continuous blood flow monitoring and laying a technical foundation for the development of truly wearable physiological monitoring devices.

However, this study still has some limitations. First, the experimental environment involves numerous instruments, introducing significant environmental noise interference, and human signals are more complex, resulting in the inability to effectively extract frequency shift information in current human experiments. Second, data acquisition still relies on large electronic equipment and has not achieved full wearability. To address these shortcomings, future research directions will focus on optimizing the patch structure to improve anti-interference capability, promoting the implementation of human experiments, and simultaneously achieving true wearable design of the device through high integration and miniaturization of the excitation and acquisition circuits, further enhancing the clinical translation value of this technology.

## Acknowledgements

This work was supported in part by the National Natural Science Foundation of China (Grant No. 12204307).

## Contributions

**Yuanlong Li:** Writing – original draft, Investigation, Data curation. **Ziqi Li:** Methodology, Visualization, Validation. **Chang Peng:** Writing – review & editing, Funding acquisition, Supervision, Conceptualization, Project administration.

